# Clustered transposon insertion via formation of chromatin loops

**DOI:** 10.1101/2022.02.16.480760

**Authors:** Roshan Prizak, Lennart Hilbert

## Abstract

Transposons, which are DNA sequences that can move to new positions in the genome, make up a large fraction of eukaryotic genomes and occur in clusters. The insertion of transposons into the genome is hindered by compact folding of chromatin, supposedly preventing aberrant or even pathogenic insertion. Chromatin can, however, be decompacted as a consequence of transposon insertion, leading to increased accessibility and, in consequence, further insertions. While these observations suggest a positive feedback between chromatin unfolding and transposon insertion, how such a feedback might contribute to clustered transposon insertion remains poorly understood. In this study, we analyze polymer models of a self-interacting chromatin domain that unfolds as increasing numbers of transposons are inserted and block the self-interaction. On the one hand, we find that, if additional transposons are inserted adjacently to already inserted transposons, the unfolding of the chromatin domain changes from a sharp globule-coil transition to a more gradual extension of loops from a core that remains folded. On the other hand, we find that adjacent transposon insertion emerges either when transposases are excluded from densely packed chromatin, or when transposon insertion proceeds very quickly in relation to the thermal equilibration of polymer configurations. We thus derive from our model physical conditions for clustered transposon insertion and the resulting spatial compartmentalization of chromatin. An according role was recently suggested for LINE-1 and Alu repeats, which occur in clusters and drive the mesoscopic compartmentalization of the mammalian genome.

**Significance Statement:** A large part of the genome is composed of repetitive sequences, so-called transposons. Transposons are involved in important processes, such as early embryonic development or control over which genes are used by the cell. Transposons frequently occur in clusters, where many similar sequence motifs are grouped together. Recent studies suggest that the insertion of transposons can result in local unfolding of the genome, favoring insertion of yet more transposons. Our work simulates a simplified region of the genome and transposases, which are the molecules that insert transposons into the genome. Surprisingly, large and fast-acting transposases favor the formation of distinct loops that contain most of the inserted transposons, providing a potential explanation for the clustered insertion of transposons.

---

**T**ransposons, also called transposable elements, are sequences of DNA that are capable of inserting themselves into new locations in the genome. Transposon insertion often disrupts the function of existing DNA sequences, such as regulatory sequences and genes, and can compromise genome stability (1). These disruptions can lead to disease (2), including carcinogenesis and tumor progression (3–5). Accordingly, organisms have developed mechanisms to prevent transposition and remove or repress transposons (6, 7). However, due to high transposition rates, transposons also contribute to genotypic and phenotypic variation (8–10). As major drivers of genome evolution, they often get retained on the genome and acquire new functions. For instance, they contribute to gene regulation (11), replication (1) and development (12). Transposons make up a surprisingly large fraction of eukaryotic genomes – about 46% of the human genome, and an astonishing 85% of the maize genome, highlighting their ubiquity.

Bioinformatic analysis of reference genomes reveals that transposons are not uniformly distributed throughout the genome, but in clusters. This transposon distribution is shaped by a combination of insertion processes and evolutionary changes in DNA sequence, but the relative contributions of these processes is still under debate (13). De novo insertions cluster on the 10 kb - 1 Mb scale, and often show a preference for regions containing active genes (13, 14). Transposon insertion is mediated by biological molecules like transposases and ribonucleoprotein complexes, which have to navigate the nuclear space, shaped in part by the three-dimensional (3D) organization of chromatin (15–19). The transposase-accessibility of chromatin can be assessed by the according ATAC method (20). Combining this method with sequencing (ATAC-seq) reveals differences in accessibility at the level of the chromatin fibre and nucleosome arrays (20). Combination with fluorescent tagging and imaging (ATAC-see) suggests that also at scales of 100 nm and above, decompacted chromatin is more accessible to transposases (21). Differential accessibility for transposition might result from steric restriction (15, 17, 19, 22) or obstruction of the formation of relevant macromolecular assemblies by the surrounding chromatin mesh-work (23). Accordingly, differences in large-scale chromatin organization seem to play a role in the non-uniform insertion of transposons.

Recent studies have pointed out how transposon activity could, in turn, shape 3D chromatin organization. LINE-1 and Alu repeats, for example, have been implicated in the spatial separation of the genome into the A and the B compartment (24). Transposons can also serve as contact domain boundaries, as illustrated by the MERVL family in the early mouse embryo (25), the HERV-H family in human pluripotent stem cells (26), and the MIR family in CD4^+^ T cells (27). Transposon insertion can also perturb chromatin organization, for instance, by disrupting sequence elements that set up chromatin organization, and lead to increase in chromatin accessibility (12, 25). Transposon activation and transcription can also induce chromatin unfolding and relocation of transcribed regions into the active nuclear compartment, away from inactive chromatin (28–36). Thus, both transposon insertion and transposon activity can trigger chromatin unfolding, potentially leading to chromatin configurations that favour transposase accessibility.

These observations suggest a possible feedback between transposition-mediated chromatin unfolding and further propensity for transposon insertion. While such a feedback can be expected to favor clustered insertion, it is still poorly understood under which conditions and by what exact process this feedback might take effect. In this study, we used polymer simulations to assess how the positive feedback between transposon insertion and chromatin unfolding affects transposon distribution. We find that unclustered insertion of transposons leads to a sharp unfolding of chromatin, while clustered insertion leads to a gradual unfolding in the form of loops. Further, we find that larger transposases and faster transposition favor this unfolding in the form of loops, resulting in clustered transposon insertion.

## Model overview

We model a 300-kb region of chromatin as a polymer chain with *N*_tot_ = 100 monomers. Monomers can be of two types - native or transposed. Only native monomers can cross-link with each other when they are in close proximity (Fig. 1a). Transposon insertion converts a native monomer to a transposed monomer, which cannot cross-link with other monomers. Each monomer represents a 3-kb chromatin region, and is assumed to have an effective diameter of 30 nm. We assume a persistence length of 3 monomers, or 90 nm, in line with experimental estimates of 50-200 nm (37), and consider harmonic bonds between adjacent monomers of the linear polymer chain. The cross-linking energy of native monomers is 1.5 *k_B_T*, in line with parameters used in previous chromatin polymer simulations (37, 38). All parameters are assigned effective values that include the influence of typical chromatin-associated proteins and RNAs.

**Fig. 1.**
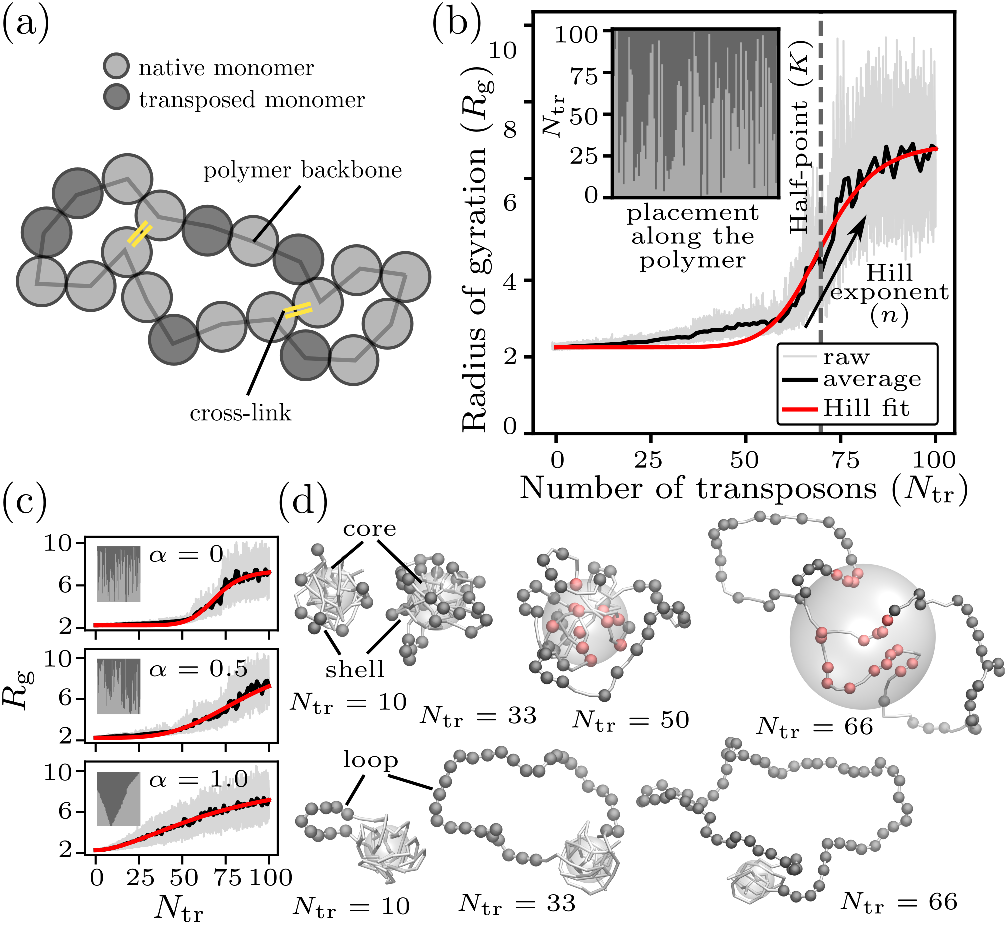
Adjacent transposon insertion converts a sharp unfolding transition into gradual loop formation. (a) Schematic of the model chromatin polymer consisting of native monomers, which can form cross-links (yellow) with each other, and transposed monomers, which cannot form such cross-links. (b) The radius of gyration (*R*_g_) of a chromatin polymer of fixed length (*N*_tot_ = 100) increases as transposons accumulate. Transposons accumulate one at a time, starting from a polymer with no transposons. The insertion history, referring to the sequence of positions on the linear polymer where transposons are inserted, is depicted in the inset. We fit a Hill curve to *R*_g_ (*N*_tr_) to capture the sharpness of unfolding and the transition point. (c) A placement adjacency (*α*) of 0 leads to sharp unfolding, non-zero values of *α* lead to gradual unfolding. (d) Example snapshots (top row: *α* = 0, bottom row: *α* = 1) from the polymer simulations at different numbers of transposons (*N*_tr_). Native monomers are shown as a white backbone. A semi-transparent white sphere with a radius equal to 40% of the bounding sphere for native monomers is shown to depict the polymer core. Transposed monomers are shown either in dark gray (if they are outside the semi-transparent sphere), or in pink (if they are inside the semi-transparent sphere).

Each simulation run starts from a compacted polymer configuration with only native monomers, and proceeds by alternating between transposon insertions and simulation segments during which polymer dynamics are evaluated (Fig. 1b). Transposon insertions are implemented differently in two different versions of the model. In the “basic model”, a placement adjaceny parameter, *α* ∈ [0, 1], is fixed at the beginning of each simulation run. For each insertion, we choose an adjacent insertion position with probability *α*. At *α* = 0, we generate polymers with insertion positions randomly chosen from all available positions, and at *α* = 1, we generate polymers with insertion positions picked only from available positions that are adjacent to existing transposons (Fig. 1c). All simulation segments are of equal duration, containing enough timesteps to allow the polymer to equilibrate between consecutive insertions. For each simulation run and number of inserted transposons *N*_tr_, we quantify the overall level of compaction by computing the polymer’s radius of gyration, *R*_g_ (Fig. 1b,c). In the “extended model”, the interaction of transposases with chromatin is modelled explicity. A transposase molecule that is near a native monomer can convert it into a transposed monomer, marking a successful transposition. The transposase is then removed from the simulation, and a new transposase molecule is added at the boundary of the simulation volume to conserve the total number of transposases.

## Results

### The chromatin polymer unfolds as the number of transposons increases

How does an increasing number of transposons affect the chromatin polymer configuration? Chromatin polymers with few transposons adopt a compact configuration facilitated by cross-links between the native monomers. With further transposition, the polymer unfolds into an open configuration. As the polymer’s radius of gyration *R*_g_ represents the effective size of the polymer, this unfolding is quantified by the observed increase in *R*_g_ with the number of transposed monomers *N*_tr_ (Fig. 1b). The unfolding proceeds similar to the well-known coil-globule (CG) transition (SI Fig. 1) (39). Here, the number of transposons at which the CG transition occurs (transition point) depends on the cross-linking energy, with weaker cross-linking leading to unfolding at a lower number of transposons. We fit a Hill curve to *R*_g_(*N*_tr_) to capture the sharpness of unfolding in terms of the Hill exponent *n* and the transition point in terms of the half-value *K* (Fig. 1b; see Appendix for details). A larger Hill exponent characterizes sharper unfolding, and a larger half-point characterizes a transition at a higher number of transposons.

### Clustered insertion of transposons leads to gradual unfolding

We can now use our simulations to assess how clustered transposon insertion might affect chromatin unfolding. To this end, we perform transposon insertion for different placement adjacency values *α*, and assess the resulting insertion histories and unfolding trajectories. By varying *α* ∈ [0, 1], we obtain insertion histories ranging from no placement adjacency (*α* = 0) and hence no clustering, to maximal placement adjacency (*α* = 1) resulting in complete clustering (insets, Fig. 1c). Placement adjacency *α* strongly affects the CG transition – while *α* = 0 results in a sharp, switch-like unfolding similar to the canonical CG transition, transposon clustering at *α* = 1 results in a gradual unfolding. In other words, when transposons are prefererentially inserted in transposon-adjacent positions, transposon clusters emerge and the polymer unfolding becomes more gradual (Fig. 1c).

The influence of placement adjacency on polymer unfolding upon *N*_tr_ can be understood by visualizing the 3D polymer configurations (Fig. 1d). As the first transposons get inserted, they form a shell on the outer layer of the compact polymer core. For low *α*, as transposons accumulate further, the shell grows, but no unfolding occurs as long as the polymer core is intact (core-shell configuration). The core starts to open up around the transition point, leading to a sharp unfolding. On the other hand, clustered transposon insertion at large *α* leads to a transposon cluster looping away from the compact polymer core (loop configuration). This loop increases in size with further transposition, leading to a gradual increase in *R*_g_. Taken together, we find that while unclustered transposon insertion leads to sharp unfolding via the core-shell polymer configuration, clustered transposon insertion leads to gradual unfolding via loop formation.

### Larger transposases and rapid transposon insertion favor clustered insertion

The observation that clustered transposon insertion leads to gradual unfolding begs the question of how such an insertion history might emerge. Transposons are preferentially inserted in less densely packed chromatin (19, 21), suggesting that clustered insertion might arise from chromatin packing differences. To further investigate this possible mechanism, we now analyze our extended model, in which we explicitly treat transposase molecules (Fig. 2a-b). In the basic model considered before, the position of each insertion, and consequently the insertion history, depend on an assumed value of placement adjacency *α*. In the current model, insertion history depends on the access of transposase to different parts of the polymer chain and emerges out of the transposase-chromatin interactions in 3D space. We assess the impact of two properties of the transposases. First, we consider size-based exclusion of transposase molecules from dense chromatin by varying transposase size, *s_T_* (Fig. 2b). Exclusion of macromolecules from densely folded chromatin has been theoretically discussed (15, 17) and experimentally shown (16, 22). Effective pore sizes of 16-20 nm were determined for highly compacted chromatin (16, 40), matching well with the diameter of proteins ranging from 5-20 nm (17), implying the possibility of volume-based exclusion of large proteins from highly compacted chromatin. Second, we vary the rate of transposition, *k_T_* (Fig. 2a). While the spontaneous rates of transposition are low (smaller than 1 per genome per generation), transposition bursts with high rates of transposition (5-10 transpositions per genome per generation) have been shown to occur in stress-related conditions in various species (10, 41, 42). We consider a wide-range of transposition rates, with low rates accounting for transpositions slower than, and high rates accounting for transpositions much faster than the chromatin polymer equilibration time.

**Fig. 2.**
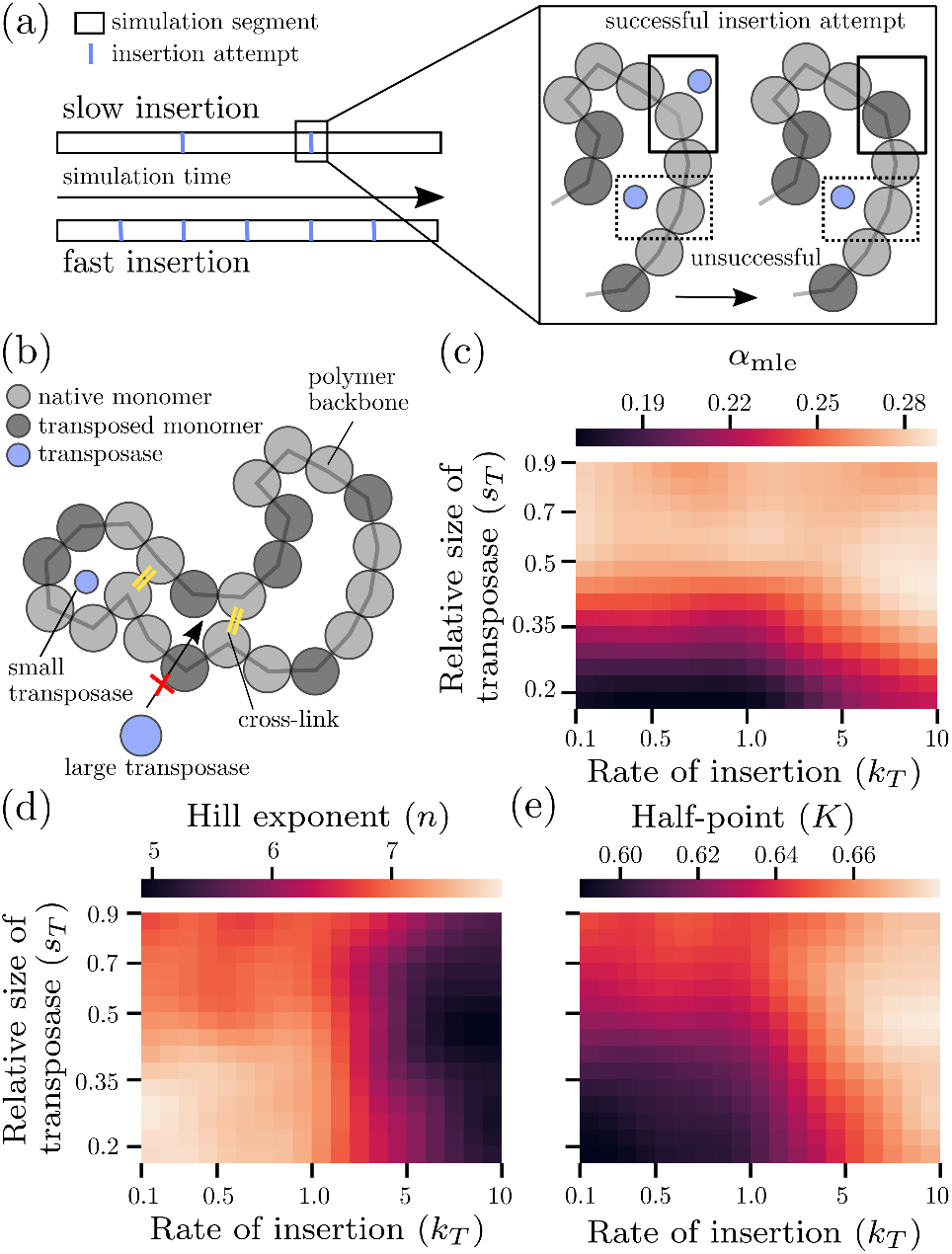
Large transposases and fast insertion result in gradual unfolding and adjacent transposon insertion. (a) A simulation run is composed of multiple simulation segments interspersed by transposon insertion attempts. In each simulation segment, the chromatin polymer and other molecules in the system move according to the potentials they experience. In each insertion attempt, native monomers close to a transposase are converted to a transposed monomer. The transposon insertion rate is implemented by changing the number of iteration steps in the simulation segments in between consecutive insertion attempts. (b) Schematic of the polymer model with transposon insertion mediated by diffusing transposases. Differential accessibility of chromatin is considered via a size-based exclusion, which restricts access to compacted polymer regions. (c) The maximum-likelihood estimate *α*_mle_ for simulation runs for different values for the rate of insertion *k_T_* and relative size of transposase *s_T_*. (d-e) Hill exponent *n* and half-point *K* of the Hill curve fitted to *R*_g_(*N*_tr_) for different values of *k_T_* and *s_T_*.

To characterize the polymer unfolding profile, we fit Hill curves to *R*_g_ (*N*_tr_) traces for different transposase sizes and insertion rates (Fig. 2d-e). For large transposases or fast insertion, the resulting Hill exponents are small, pointing to gradual unfolding as observed previously for large *α*. On the other hand, for small transposases or slow insertion, the resulting Hill exponents are larger, pointing to sharp unfolding as observed previously for small *α*. In the basic model, we saw that *α* and the resulting transposon clustering determined the unfolding profile. To check whether the differences in Hill exponents result from differences in transposon insertion bias also in the extended model, for each insertion history, we perform a maximum likelihood estimation of the underlying placement adjacency *α*_mle_ (Fig. 2c, see Methods and Appendix for details). We find that smaller transposases and slower insertion result in fewer adjacent insertions, consistent with the sharp unfolding and large Hill exponents. Thus, both larger transposases and quicker insertion result in more adjacent insertions, in line with gradual unfolding and small Hill exponents (Fig. 2d-e).

### Large transposases and high insertion rates favor transposition in the polymer shell

Based on the observation of a core of native monomers in some of our simulations, we hypothesized that the ability of transposases to access this core is related to clustered transposon insertion. Indeed, for large transposases and high insertion rates, the mean normalized insertion distance was increased (Fig. 3a). A more detailed analysis confirms that fast insertion and large transposase size lead to more insertion within the shell and less insertion in the polymer core (Fig. 3b). When fast insertion and large transposase size are combined, insertion is almost fully restricted to the outer shell (Fig. 3b). This restriction of insertions towards the outer shell is reflected by an exclusion of transposases from the polymer core (Fig. 3c).

**Fig. 3.**
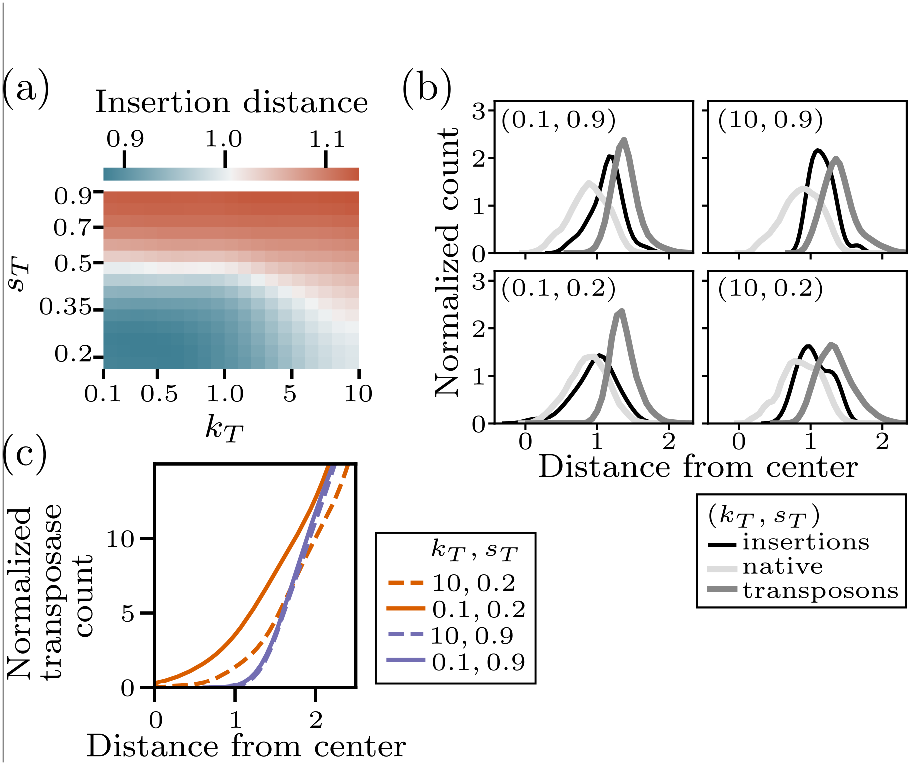
Clustered insertion is driven by exclusion of transposases from the polymer core. (a) Average insertion distance (first 30 insertions, relative to polymer center, normalized by *R*_g_) for different transposase size *s_T_* and insertion rate *k_T_*. (b) Distance distribution native monomers, transposed monomers, and insertion events (first 30 insertions) for four different sets of *s_T_* and *k_T_*. (c) Distance distribution of transposases for four different sets of *s_T_* and *k_T_*.

### Clustered transposon insertion is associated with unfolding via loop formation

Taken together, our model analysis suggests four scenarios how chromatin unfolding relates to progressive transposon invasion (Fig. 4a). Scenario I, which occurs for small transposases and slow insertion, is characterized by polymer unfolding via the core-shell mechanism (Fig. 4b). Scenarios II and III are characterized by unfolding via several small loops (Fig. 4b). In scenario II, large transposases cannot penetrate past the outer polymer shell, leading to clustered transposition. In scenario III, small transposases can, in principle, access the polymer core, but due to their fast transposition rate become exhausted before actually reaching the core. In scenario IV, in which transposons are large and insert transposons quickly, typically only one prominent loop emerges (Fig. 4b).

**Fig. 4.**
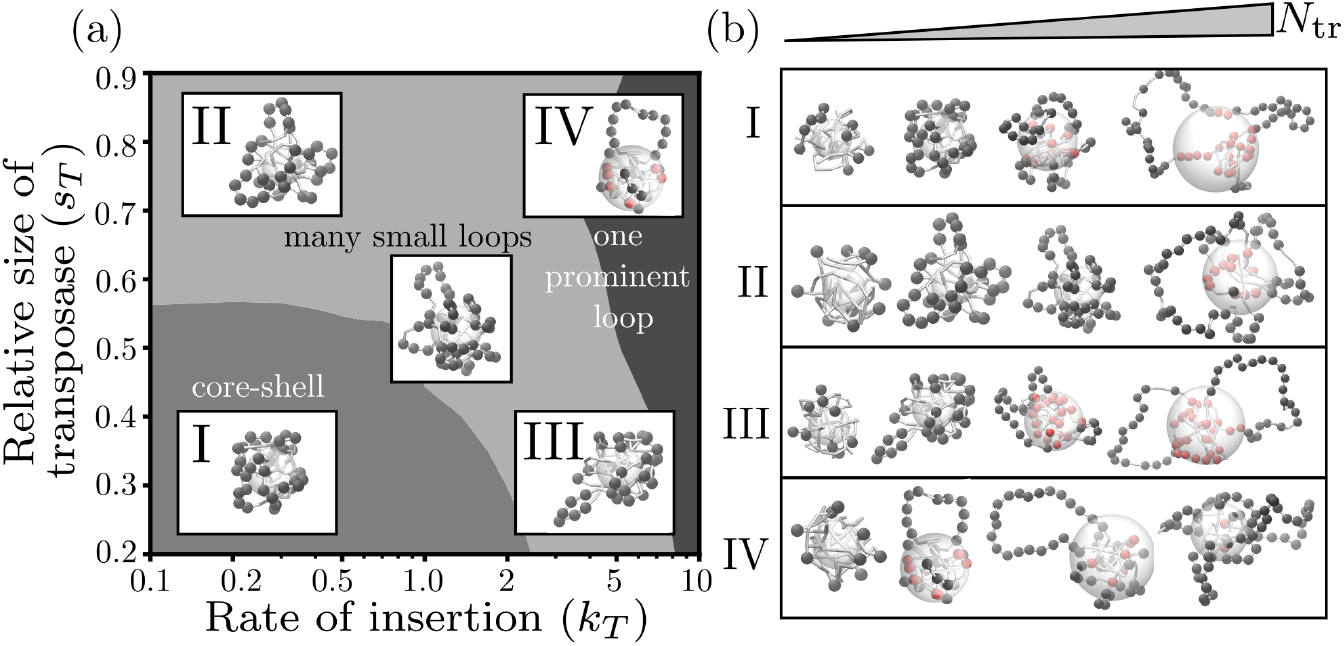
Different scenarios of transposon insertion and polymer unfolding. (a) Modulation of transposase size and insertion rate leads to four different scenarios of transposon-induced unfolding. In scenario I, small transposases and slow insertion lead to unclustered insertion and polymer configurations consisting of a compact core of native monomers and a shell of transposed monomers around it. Scenarios II and III exhibit unfolding via the formation of several small loops that occur as a result of clustered insertion - either because of size-based exclusion of large transposases or fast insertion on the outer shell. Scenario IV typically exhibits one prominent loop of clustered transposons. (b) Representative snapshots of polymer simulations at increasing numbers of transposons in the four scenarios (left to right, *N*_tr_ ≈ 10, 33, 50, 66).

## Discussion

In this study, we used polymer simulations of a model chromatin domain to study how clustered transposon insertion can arise from a feedback between transposon insertion and chromatin unfolding. We found that large transposons as well as fast transposon insertion favor clustered insertion. The crucial assumption for this effect is that transposon integration drives unfolding of chromatin, leading to the preferred integration of further transposons at loops that protrude from a globular polymer core formed from chromatin without inserted transposons. Indeed, in polymer models of retroviral insertion, insertion in decompacted is highly favored (19). These findings differ from the intuitive expectation that smaller molecules should access chromatin more easily, and faster insertion should favor transposon invasion. While our model is a strong simplification of the actual transposition process, it still presents an explanation of how changes in chromatin folding can shape the course of transposon invasion. Clustered transposon insertion, as predicted in our model, has been suggested as a major driver of the three-dimensional compartmentalization of chromatin regions with different epigenetic states. For example, transcriptional activation of non-coding LINE-1 repeat elements after fertilization regulates global chromatin accessibility in the early mouse embryo (12). Also, the non-coding repeats LINE-1 (clustered within heterochromatin regions) and Alu (clustered within euchromatin regions) exhibit homotypic interactions, leading to formation of the two spatially separated genomic A and B compartments (24, 43). Lastly, the relatively unfolded state of euchromatin has been associated with the SAF-A (also called U1 snRNP or HNRNPU) mediated retention of coding as well as non-coding RNAs (35, 44, 45). Many of the retained RNAs are transcribed from clustered non-coding elements (46), and our work provides a model for the emergence of such clustered placement of non-coding elements.

We assume chromatin compaction via cross-links and size-based transposase exclusion as key mechanisms underlying the clustered insertion of transposons. Other mechanisms have also been suggested as causes for clustered transposon insertion. First, considering chromatin compaction and unfolding, several relevant biophysical mechanisms have been identified (47, 48). For example, heterochromatin can be compacted by embedding in liquid-like or gel-like membrane-less organelles (49, 50). Another mechanism that was suggested for heterochromatin compaction is a polymer collapse transition, similar to the coil-globule transition seen in our model (51). A model that was recently applied to the formation of compacted chromatin clusters by the Structural Maintenance of Chromosomes (SMC) proteins is bridging-induced attraction (38, 52). Unfolding of euchromatin, and specifically regions harboring transcriptionally active genes, was explained via a dispersing effect of nascent and chromatin-associated RNA transcripts, leading to microphase separation from inactive chromatin (35, 36). Second, considering the aspect of feedback between chromatin modifications to transposon insertion, physical mechanisms different from chromatin decompaction have also been investigated in terms of their effect on chromatin accessibility. For example, a screen of several hundred transcription factors revealed that, whereas some of the DNA-binding factors are excluded from compact chromatin, others have a strong bias to colocalize with chromatin without being affected by compaction (18). Another mechanism that might limit DNA modification to distinct domains is the formation of microphase-separated contact domains that restrict the reach of chromatin-binding enzymes (53–55). Formation of such contact domains was attributed to the microphase separation of epigenetically distinct chromatin regions and was also described by block co-polymer simulations (56–58). Another relevant physical property of the chromatin fibre might be the local stiffness, with previous work proposing that genomic loci with lower bending stiffness might favor the integration of transposons (59). Lastly, super-coiling of the chromatin fibre was found to correlate with transcriptional activity and chromatin decompaction (60). Polymer simulations show that super-coiling can lead to clustered insertion via a positive feedback loop similar to the positive feedback we found for chromatin decompaction (61). From these different mechanisms and explanations, a common theme emerges: clustered insertion occurs via a positive feedback between a chromatin modification resulting from transposon insertion and further, localized insertion of transposons. Different processes can instantiate this type of feedback, implying that clustered transposon insertion can emerge on the basis of several biophysical mechanisms.

In our model, transposases diffuse inward from the walls of the volume containing the simulated 300-kb chromatin polymer. This picture matches the “copy-and-paste” transposition mechanism. However, except for insertions occurring within this local chromatin domain, our model would be the same for “cut-and-paste” transposition. A polymer of 300 kb length also matches the scale at which clustered de novo insertions of LINE-1 transposons was observed (13). Chromatin domains of this size hierarchically associate into larger compartment domains (54, 62), raising the question of how transposon insertion within a given 300-kb domain is related to this overall genome organization. Similar to recent findings on non-homologous double-strand break repair, the spreading of transposon insertion might be limited by a CTCF-mediated, relative reduction of 3D chromatin-chromatin contacts between neighboring domains (55). A better understanding of how changes in chromatin accessibility might proliferate into higher-order 3D interactions would require the deployment of multi-scale polymer models, providing a relevant direction for future work.

An uncontrolled sequence of transpositions, called transposon invasion, can severely disrupt genome integrity. Metazoans and plants have developed mechanisms to protect against transposon invasions, such as the piRNA pathway (63, 64) or specific siRNAs (65). Our work suggests that certain modes of transposon insertion that lead to gradual unfolding of chromatin could make the affected chromatin region vulnerable to further invasion. Perhaps surprisingly, small transposases, which lead to sharp unfolding, could in fact be less effective in an invasion, as their unbiased insertion reduces chromatin unfolding during the initial transposition events. Transposon insertion and removal processes span a wide range of timescales, starting from the physiological timescale, on which transposon insertion and transposon silencing occur, via the inter-generational timescale of transposon silencing, up to evolutionary timescales of transposon removal or proliferation (66). To fully investigate the dynamics of transposon invasion, including parameter regimes or situations which offer protection against invasion, will require models that consider transposon insertion as well as transposon silencing or removal on various timescales.

## Materials and Methods

### Model details

Monomers in the chromatin polymer and transposases (referred to as atoms henceforth) interact with phenomenological force fields (potentials) and follow Netwon’s laws of motion. We simulate the spatial dynamics of the system in the molecular dynamics simulator LAMMPS (67). At the core of the simulation is a standard Velocity-Verlet algorithm that solves the underlying Langevin equations describing the system dynamics. The chromatin polymer is held together by harmonic springs between adjacent pairs of monomers, and the polymer’s bending rigidity is modeled by a Kratky-Porod potential. To model pure (steric) repulsion between any pair of atoms, we consider the Weeks-Chandler Andersen (WCA) potential, which is a shifted version of the Lennard-Jones (LJ) potential with a cutoff at zero-crossing point. To model steric repulsion, this cut-off distance is typically set to be the mean diameter of the two interacting atoms. As all the monomers have a diameter *σ* = 30 nm, for all monomer pairs, the cutoff distance is set as *r*_cut_ = 2^1/6^*σ*. Transposases have a size *s_T_σ*, so the cutoff distance for the repulsive WCA potential between any pair of transposases is 2^1/6^*s_T_σ*. For a monomer-transposase pair, we set the cutoff distance to be 2^1/6^*s_T_σ*, a value smaller than the mean diameter of a monomer-transposase pair. This allows the transposase molecule to come closer to a monomer molecule than another monomer molecule can. Moreover, the cutoff distance is smaller for smaller transposases, allowing them to come closer to monomers than larger transposases. In effect, this models size-based exclusion of transposases from chromatin. Cross-linking interactions (crosslinking energy *E* = 1.5 *k_B_T*) are modeled by using a shifted and truncated Lennard-Jones potential with a larger cut-off distance of 2.5*σ*. If a pair of native monomers are within a distance between 2^1/6^*σ* and 2.5*σ*, they experience an attraction and form a cross-link.

### Simulation run

Each simulation run in LAMMPS starts with an initial polymer with no transposons, and consists of an initial “soft” phase, an equilibration phase, and finally the main simulation block in which transposons are sequentially inserted into the chromatin polymer. In the basic model without transposases, the main simulation block contains 100000(*N*_tot_ + 1) time-steps, insertion is executed every 100000 time-steps. In the transposase model, insertion is attempted every 1000*/k_T_* time-steps, leading to 100 – 10000 time-steps per simulation segment. In between simulation segments, every native monomer that has a transposase within a distance of *σ*(1 + *s_T_*)/2 is converted to a transposed monomer, and the transposase is moved to the simulation volume boundary.

### Placement adjacency

If a native monomer has a transposon adjacent to it, depending on the value of the placement adjacency *α*, it can preferentially get the next transposon as compared to a native monomer without any adjacent transposon. At *α* = 0, all native monomers are have equal insertion probability, independent of being adjacent to transposons or not. At *α* = 1, only native monomers adjacent to transposons receive further insertions. In the transposase model, we perform a maximum likelihood estimation of the underlying *α* as *α*_mle_ by using the first 50 insertions from multiple independent repeat runs.

## ACKNOWLEDGMENTS

This work was supported by the Helmholtz Program “Natural, Artificial, and Cognitive Information Processsing”. The study was started by discussions with Changjing Zhuge and Jinzhi Lei during an exchange visit supported by the Sino-German Center for Research Promotion. Davide Michieletto advised on the implementation of conversion reactions in LAMMPS simulations and commented on the manuscript. The authors declare that they have no conflict of interest.

