## Supplementary Information & Materials for "Clustered transposon insertion via formation of chromatin loops"

#### This PDF file includes:

Supplementary text
Fig. S1
SI References

### Supporting Information Text

All atoms in the system (monomers and transposases) interact with phenomenological force fields (potentials) and follow Netwon's laws of motion. The following Langevin equation describes the spatial dynamics of the $i^{\text{th}}$ atom:

$$m_i \frac{\mathrm{d}^2 \mathbf{r}_i}{\mathrm{d}t^2} = -\nabla U_i - \gamma_i \frac{\mathrm{d}\mathbf{r}_i}{\mathrm{d}t} + \sqrt{2k_{\mathrm{B}}T\gamma_i}\eta_i(t), \quad [1]$$

where $\mathbf{r}_i$ is the position of $i^{\text{th}}$ atom, $m_i$ its mass, $\gamma_i$ the friction on it from the solvent, $U_i$ the potential experienced by it, and $\eta_i$ is random uncorrelated noise with zero mean. A standard Velocity-Verlet algorithm is used to solve Eq. 1 in LAMMPS. The potential $U_i$ is formed by accounting for all interactions between the $i^{\text{th}}$ atom and all other atoms. The chromatin polymer is made up of $N_{\text{tot}}$ monomer sub-units, formed by harmonic springs between every pair, $i$ and $i+1$, of adjacent monomers with $K_{\text{harmonic}} = 90k_{\mathrm{B}}T$ as:

$$U_{\text{harmonic}}(r_{i,i+1}) = K_{\text{harmonic}}(r_{i,i+1} - R_0)^2, \quad [2]$$

where $r_{i,i+1} = ||\mathbf{r}_i - \mathbf{r}_{i+1}||$ is the distance between adjacent monomers. We also consider a Kratky-Porod potential to model the bending rigidity of the polymer

$$U_{\text{BEND}}(\theta) = K_{\text{BEND}}[1 - \cos\theta], \quad [3]$$

where $\theta$ is the angle between every three adjacent monomers, and $K_{\text{BEND}}$ is the bending energy. Persistence length is given by $L_p = K_{\text{BEND}}\sigma$. Repulsion between atoms is modeled as a Weeks-Chandler-Andersen (WCA) potential, which is a shifted and truncated version of a Lennard-Jones (LJ) potential. The WCA potential for a pair of atoms $i$ and $j$ is

$$U_{\text{WCA}}(r_{ij}) = \begin{cases} 4\left[\left(\frac{d_{ij}}{r_{ij}}\right)^{12} - \left(\frac{d_{ij}}{r_{ij}}\right)^{6}\right] + 1, \text{if } r_{ij} < 2^{\frac{1}{6}} d_{ij} \\ 0, \text{otherwise,} \end{cases} \quad [4]$$

with $r_{i,i+1} = ||\mathbf{r}_i - \mathbf{r}_{i+1}||$ is the distance between the atoms, and $d_{ij}$ is usually the mean diameter of the atoms $i$ and $j$ (See below for details on size-based exclusion).

Monomers come from one of two types, native and transposed. Native monomers can cross-link among themselves with a cross-linking energy $E$, while transposed monomers cannot form cross-links. Native monomers represent the default chromatin monomer type while transposed monomers represent those with a transposon insertion and hence cannot form cross-links. These cross-linking interactions are modeled as shifted, truncated Lennard-Jones potential

$$U_{\text{LJcut}} = \begin{cases} U_{\text{LJ}_0}(r_{ij}) - U_{\text{LJ}_0}(r_{\text{cut}}), \text{if } r_{ij} < r_{\text{cut}} \\ 0, \text{otherwise,} \end{cases} \quad [5]$$

where the Lennard-Jones potential $U_{\text{LJ}_0}(r)$ is

$$U_{\text{LJ0}}(r) = 4E\left[\left(\frac{d_{ij}}{r}\right)^{12} - \left(\frac{d_{ij}}{r}\right)^{6}\right], \quad [6]$$

and the cut-off distance is given by $r_{\text{cut}} = 2.5\sigma$. If cross-linking monomers are within a distance between $2^{1/6}\sigma$ and $r_{\text{cut}}$, they feel an attraction.

**Simulation parameters.** In the chromatin polymer, each monomer corresponds to a chromatin segment of size 3 kbp and has a diameter $\sigma = 30$ nm, assuming a compaction of around 100 bp/nm. We consider $K_{\text{BEND}} = 3\ k_{\mathrm{B}}T$, resulting in $L_p = 3\sigma = 90$ nm. The persistence length of chromatin is not understood well, but experimentally observed values range between 50-200 nm (1). We consider a default cross-linking energy of $E = 1.5\,k_{\mathrm{B}}T$. Each run of a polymer simulation consists of 1 million timesteps, with each timestep of length $dt = 0.01\tau$, where $\tau = \sqrt{\sigma^2 m/k_{\mathrm{B}}T}$ is the natural timescale of the system. All components of the system are assumed to have the same mass ($m = 1$).

In the transposase model, we have 25 transposase molecules that can mediate transposon insertion. We vary the transposase size $s_T\sigma$ between $0.2\sigma = 6$ nm and $0.9\sigma = 27$ nm, and the insertion rate over two orders of magnitude $k_T \in [0.1, 10]$, corresponding to a fastest rate of an insertion attempt every 100 timesteps and a slowest rate of an insertion attempt every 10000 timesteps.

**Simulation steps.** Each simulation run consists of three phases - an initial "soft" phase, an equilibration phase, and then the main simulation phase.

**Insertion process.** The chromatin polymer with a total of $N_{\mathrm{tot}}$ monomers and $N_{\mathrm{tr}}$ transposons inside it can be represented as $s$, a binary string of length $N_{\mathrm{tot}}$ with 0's representing native monomers and 1's representing transposed monomers. As the chromatin polymer is joined at the ends, any shifted version of $s$ is equivalent to $s$. We call this the polymer state. For example, if $N_{\mathrm{tot}} = 6$, a possible polymer state with two transposons is $s(2) = 011000$. This is equivalent to its shifted versions 110000, 100001, 000011, 000110 and 001100.

A transposon insertion event consists of selecting one of the $N_{\mathrm{tot}} - N_{\mathrm{tr}}$ 0's from the current polymer state $s(2)$ and replacing it with a 1 to obtain a new polymer state $s(3)$. For example, an insertion in the first position of $s(2) = \mathbf{0}11000$ would lead to $s(3) = 111000$. However, an insertion could have occurred in one of the other three available positions (0's), giving rise to $s(3) = 011\mathbf{1}00$, $0110\mathbf{1}0$ or $01100\mathbf{1}$.

How are these insertion positions selected? Given a polymer state, each monomer $m_i$ at the $i^{th}$ position has two adjacent neighbours, usually at the $i-1$ and $i+1$ positions. The first and the last monomer in any given $s$ are neighbours of each other. Insertion can be of two types - random (R) or adjacent (A). Random insertion involves insertion into one of the available native monomers (0). Adjacent insertion involves insertion into a native monomer (0) with at least one transposon (1) as one of its two neighbours.

In the basic model, each insertion's type is first chosen from the two options - adjacent insertion type with probability $\alpha$ (placement adjacency), or a random insertion type with probability $1-\alpha$. If the chosen insertion type is adjacent, one of the available native monomers (0) which has a transposon neighbour (1) is chosen. The local substring in the polymer state around the chosen position would be one of the following three possibilities – (..1**0**0..), (..0**0**1..), or (..1**0**1..). If the chosen insertion type is random, one of the $N_{\mathrm{tot}} - N_{\mathrm{tr}}$ available native monomers (0) is picked randomly. The local substring in the polymer state around the chosen position would be one of the following four possibilities – (..0**0**0..), (..1**0**0..), (..0**0**1..), or (..1**0**1..). Note that even though the chosen insertion type is random, three of the possible chosen positions have a transposon neighbour, and hence a random insertion can also result in a seemingly adjacent insertion.

In the transposases model, insertion positions are selected based the proximity between native monomers and transposases. In an insertion event, the set of native monomers with a transposase within a distance of $(1+s_T)\sigma$ from each of their centers are selected and converted to a transposon. Unlike the basic model, the insertion positions do not depend on a pre-specified placement adjacency.

**Insertion history.** In a single simulation run, starting from a polymer with no transposons $s(0) = 000\ldots000$, transposons are sequentially inserted at various positions, with the polymer state changing with each insertion. Each insertion changes one of the 0's in the polymer state to a 1. The simulation ends when all monomers are transposed $s(N_{\mathrm{tot}}) = 111\ldots111$. The set of polymer states $s(i), i = 0, 1, \ldots N_{\mathrm{tot}}$ together is called the insertion history.

**Hill curve fit.** We fit a Hill curve to $R_{\mathrm{g}}(N_{\mathrm{tr}})$ to obtain the Hill exponent $n$ and the half-point $K$:

$$f(x) = R_{\mathrm{g}}(0) + (R_{\mathrm{g}}(N_{\mathrm{tot}}) - R_{\mathrm{g}}(0))(1+K^n)\frac{x^n}{x^n+K^n}, \qquad [7]$$

where $x = N_{\mathrm{tr}}/N_{\mathrm{tot}}$ is the fraction of transposons.

**Estimation of $\alpha$.** The insertion history in the transposase model emerges from the interactions between transposases and chromatin polymer, and the accessibility constraints resulting from transposase size and insertion rate. This is in contrast with the basic model without transposases, in which insertion history was generated from a single parameter $\alpha$, the placement adjacency. To compare the insertion histories between the basic model and the transposase model, we estimate the underlying $\alpha$ in the transposase model using a maximum likelihood estimation (MLE) method on the set of resulting histories. This MLE estimate $\alpha_{\mathrm{mle}}$ can be interpreted as the most likely placement adjacency value $\alpha$ that would generate the resulting set of insertion histories if the underlying model were the basic model.

Given an insertion history $s(i), i = 0, 1, \ldots, N_{\mathrm{tot}}$, for each insertion $s(i) \to s(i+1)$, in $s(i)$ we check if the inserted position has a transposon neighbour (..1**0**0..), (..0**0**1..), or (..1**0**1..) - say, event A, or if neither of the two neighbours are transposons (..0**0**0..) - say, event B. We also note down the total number of native monomers in $s(i)$ that have a transposon neighbour – $m$. So the number of native monomers in $s(i)$ that do not have a transposon neighbour is $N_{\mathrm{tot}} - i - m$. We define

$$p_i = \frac{m}{N_{\mathrm{tot}} - i}. \qquad [8]$$

We suppose the resulting insertion history is generated with an underlying placement adjacency $\alpha$. For each insertion, with probability $\alpha$, insertion occurs in a position that has an adjacent transposon (event A), and with probability $1-\alpha$, the insertion position is chosen randomly. However, one of the randomly chosen positions can have an adjacent transposon (probability $p_i$), and hence result in an event A. With probability $1-p_i$, an event B can occur. Hence, the overall probability of event A occurring is $\alpha + (1-\alpha)p_i$, and the probability of event B occurring is $(1-\alpha)(1-p_i)$. Each insertion in the insertion history can be classified as either an event A or an event B, leading to a string of A's and B's of length $N_{\text{tot}}$. From this, we can write the expression for the log likelihood of the first $T$ insertions of the given insertion history as

$$l(\alpha) = \sum_{i=1}^{T} \log Q_i, \qquad [9]$$

where $Q_i(\alpha) = \alpha + (1-\alpha)p_i$ if the $i^{th}$ insertion is an event A, and $Q_i(\alpha) = \alpha + (1-\alpha)p_i$ if the $i^{th}$ insertion is an event B. Now, if $r$ individual repeats of the simulation are performed, we have $r$ simulation histories $s_j(i), i = 0, 1, \ldots, N_{\text{tot}}, j = 1, 2, \ldots, r$. We can write the log likelihood of the $r$ insertion histories as

$$l(\alpha) = \frac{1}{r} \sum_{j=1}^{r} \sum_{i=1}^{T} \log Q_{ij}, \qquad [10]$$

where $Q_{ij}(\alpha) = \alpha + (1-\alpha)p_{ij}$ if the $i^{th}$ insertion of the $j^{th}$ run is an event A, and $Q_{ij}(\alpha) = \alpha + (1-\alpha)p_{ij}$ if the $i^{th}$ insertion of the $j^{th}$ run is an event B. As the likelihood $L$ is a function of the placement adjacency $\alpha$, we can find the maximum likelihood estimate of $\alpha$ as

$$\alpha_{\text{mle}} = \underset{\alpha}{\text{argmax}}\ l(\alpha), \qquad [11]$$

which is the value of $\alpha$ that maximizes the likelihood of obtaining the given $r$ insertion histories.

**Coil-globule transition.** In a lattice-framework, assuming that each lattice site has $z$ sides, the number of expected interactions between native monomers and transposons is

$$m_{AB} = zN_{\text{tr}}. \qquad [12]$$

This is different from the expected $m_{AB} = zN_{\text{tr}}(N_{\text{tot}} - N_{\text{tr}})/N_{\text{tot}}$ according to random mixing in the Bragg–Williams mean-field approximation (2). This deviation arises because for low number of transposons, the polymer is collapsed from strong native monomer interactions. For each transposon in the collapsed polymer core, the probability that it has a native monomer neighbour is 1. Continuing further (see Chapters 15 and 33 from (2)), we find that the exchange parameter

$$\chi_{AB} = \frac{1}{2} z \frac{E}{k_{\text{B}}T} \left(1 - \frac{N_{\text{tr}}}{N_{\text{tot}}}\right). \qquad [13]$$

The coil-globule transition occurs at $\chi_{AB} = 0.5$, giving us $z\frac{E}{k_{\text{B}}T}(1-K) = 1$. We estimate $z$ from the Hill curve fitted at $E = 1.5$ and use that to predict the half-points for other values of cross-linking energies (SI Fig. 1).

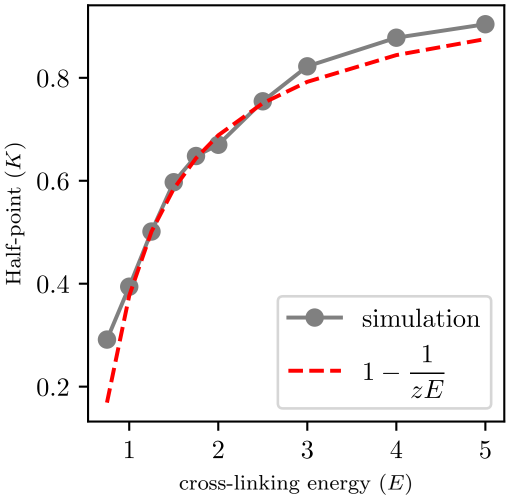


**Fig. S1. Half-point prediction according to the coil-globule transition.** We compare the half-points $(K)$ of the Hill curves $R_{\mathrm{g}}(N_{\mathrm{tr}})$ obtained from simulations and the prediction of the coil-globule transition for different values of cross-linking energies $E$.
